# Node centrality in MEG resting-state networks co-varies with neurotransmitter receptor and transporter density

**DOI:** 10.1101/2024.01.11.575176

**Authors:** Felix Siebenhühner, J. Matias Palva, Satu Palva

## Abstract

Neuronal oscillations are central mechanisms in the regulation of neuronal processing and communication (Deco et al., 2011; Fries, 2015; S. Palva & Palva, 2012; Siegel et al., 2012; Singer, 1999), but the relationship between the emergent inter-areal synchronization of oscillations and their underlying synaptic and neuromodulatory mechanisms - the dynome - has remained poorly understood (Kopell et al., 2014).

While oscillations are largely generated by fast synaptic neurotransmission among pyramidal cells and interneurons (Traub et al., 2004), these microcircuits are subject to slower neuromodulation in a frequency- and region-specific manner (Batista-Brito et al., 2018; Roopun et al., 2010). While the efferent connections, receptor densities, and neurotransmitter reuptake regulation of neuromodulatory systems are highly heterogeneous across the cortical mantle (Avery & Krichmar, 2017; Deco et al., 2017; Hansen et al., 2022), it has remained unresolved how this variability shapes inter-areal connectivity of neuronal oscillations.

Here, we used source-reconstructed human magnetoencephalography (MEG) data to assess how the centrality of brain areas (nodes) in large-scale networks of phase synchrony and amplitude correlations covaries with neurotransmitter receptor and transporter densities. Node centrality strongly covaried with receptor and transporter densities in a coupling- and frequency-specific manner both at the level of individual receptors and transporters and that of principal components.

In delta, theta, and gamma frequencies, node centrality in phase-synchronization networks covaried positively, and in high-alpha and beta bands negatively, with dopaminergic, GABA, NMDA, muscarinic, and most serotonergic receptor densities. In amplitude-correlation networks, node centrality in delta and gamma bands covaried positively, and in theta to beta bands negatively, with most receptor and transporter densities. These results establish the contribution of neurotransmitter receptor and transporter densities for shaping connectivity of neuronal oscillations in the human brain and demonstrate a link between coupling of neuronal oscillations with the underlying biological details.

## Results and Discussion

### Principal component analysis reveals neuroarchitectonical principles underlying density maps of neurotransmitter receptors and transporters

We used the neurotransmitter receptor and transporter density maps (Hansen et al., 2022) at the resolution of the 200-parcel Schaefer atlas (Schaefer et al., 2018) (Suppl. Fig. 1). Since many of these maps exhibited similar spatial patterns, we first identified the shared underlying patterns with principal component analysis (PCA). The first 5 components explained > 85% of variance in the density maps (40%, 18.1%, 12.6%, 9.1%, and 5.9%, respectively) (Fig. 1A). We then estimated the “loading” of the individual density maps to these PCs with Spearman’s correlation coefficient.

**Figure 1.**
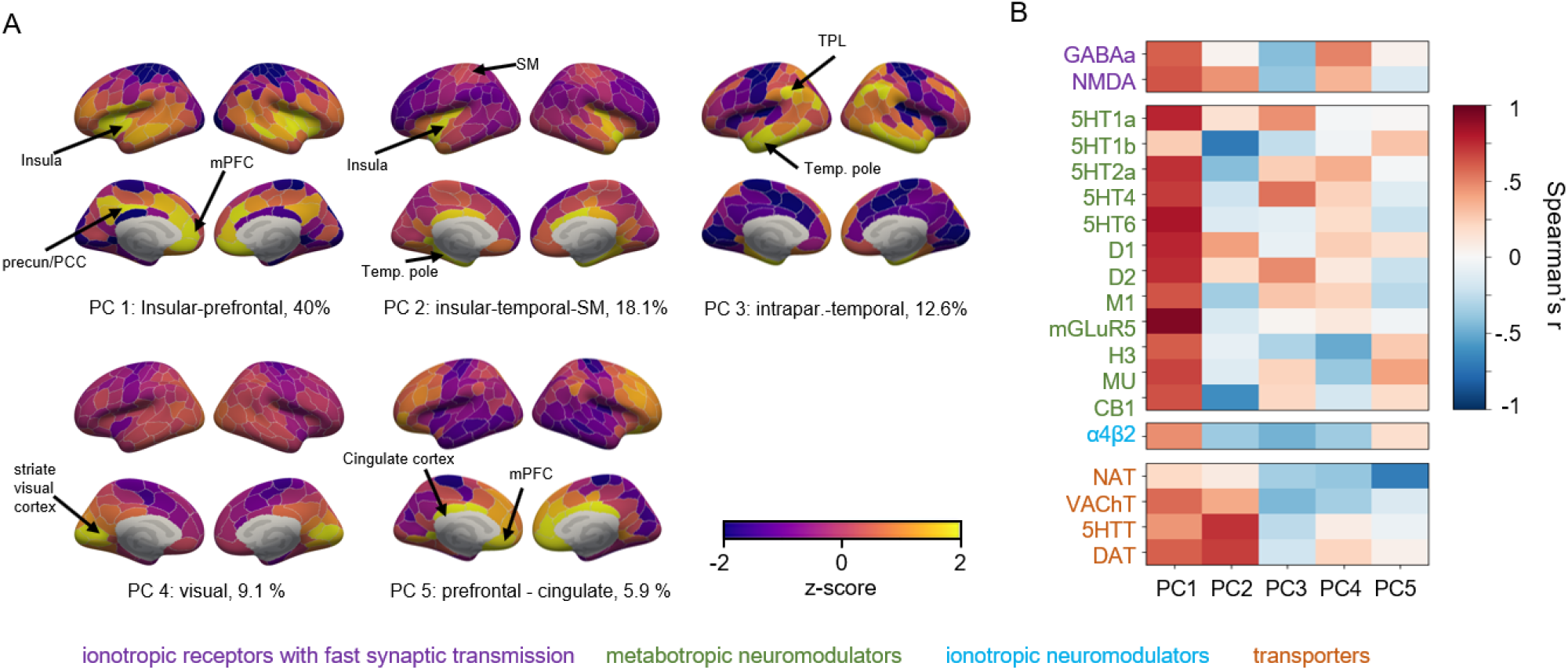
Principal component analysis reveals common spatial patterns in receptor maps. **A**: The first 5 principal components (PCs) of the receptor and transporter density maps. **B**: PC loadings of individual density maps indicate architectonical principles. **Abbreviations**: PC: principal component; PCC: posterior cingulate cortex; PFC: prefrontal cortex; SM: somatomotor cortex, IPL: intraparietal lobule; 5HT: serotonin, 5HTT: serotonine transporter, D: dopamine, DAT: dopamine transporter, M1: muscarinic ACh receptor, mGLuR5: metabotropic (G-protein-coupled) glutamatergic receptor, NAT: noradrenaline transporter, VAChT: vesicular acetylcholine transporter, a4b2: α4ß2 nicotinic ACh receptor, H3: histamine receptor 3,, MU: mu-opioid receptor, CB1: cannabinoid receptor 1.

Reproducing earlier findings (Hansen et al., 2022), the first component (PC1) was the strongest along the lateral sulcus around the insular and in medial prefrontal regions as well as the precuneus and PCC (see Fig. 1A) and was positively loaded for all density maps (Fig. 1B). PC1 thus represents an overarching neuroarchitecture of receptor and transporter distributions (Celada et al., 2013). In addition, however, we identified four more anatomically distinct principal components (PCs 2-5, see Fig. 1). PC2 was most pronounced around the insula (similar to PC1), but also in medial temporal and cingulate cortex and the only PC with positive loading on the somatomotor cortex. This component showed highest loadings with 5HTT, DAT, D1, D2, and glutamatergic NMDA and VAChT receptors and transporters. PC3, in contrast, was strongest in lateral temporal and intraparietal cortex, and had strongest loadings for 5HT1a, 5HT4, D2, MU, CB1, and M1. PC4 was largest in visual regions, and had strong loadings for serotonergic, dopaminergic, GABA, and NMDA receptors and transporters, while PC5 was strongest in prefrontal and cingulate regions and had strongest loadings for histamine, α4β2, MU, and CB1 receptors.

### Inter-areal oscillatory phase and amplitude connectivity are anatomically and spectrally specific

We performed source-reconstruction on 10-min resting-state MEG data sets from 67 healthy subjects and collapsed source time series to the 200-parcel Schaefer atlas (Siebenhühner et al., 2020). We quantified pairwise phase synchrony (PS) between parcel time series with the imaginary part of the complex phase locking value (iPLV) (J. M. Palva et al., 2018) and amplitude correlations (AC) with the orthogonalized correlation coefficient (oCC) (Brookes et al., 2011), as these metrics are less influenced by signal leakage that is a common issue in MEG (J. M. Palva et al., 2018; S. Palva & Palva, 2012) (Fig. 2A,B). In line with previous studies (Brookes et al., 2011; Siebenhühner et al., 2020; Siems & Siegel, 2020), the mean strength of iPLV and oCC exhibited a peak centered in alpha band around 10 Hz while significant phase and amplitude correlations were observable throughout the studied frequency range from delta to gamma bands (Fig. 2C).

**Figure 2.**
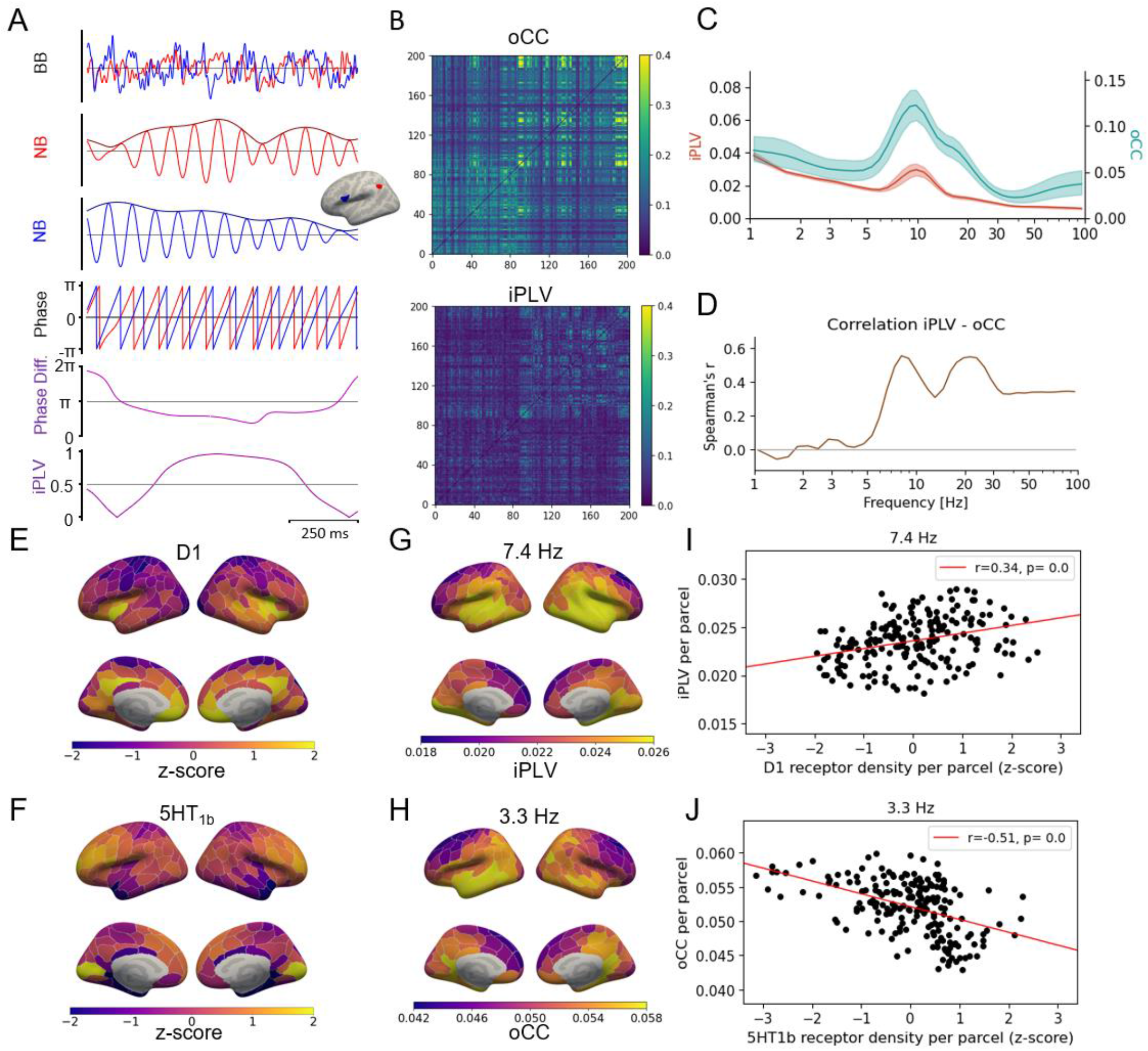
Inter-areal phase and amplitude coupling. **A**. Example broadband time series from two cortical parcels (top row), with their narrowband (10.9 Hz) real and amplitude time series (2nd, 3rd row), and phase time series (4th row) and the phase difference between them (5th row) and iPLV in sliding time windows of 364 msec length (bottom row). **B**. Amplitude correlation and phase synchrony interaction matrices, estimated with oCC and iPLV, resp., between all parcel pairs. **C**. Mean coupling strength per frequency for oCC and iPLV. **D**. Spearman’s correlation between oCC and iPLV node strength across parcels. **E**. D_1_ receptor density per cortical parcel (z-scored). **F**. Same for 5HT_1b_ receptor density. **G**. iPLV node centrality per parcel at 7.4 Hz. **H**. oCC node centrality per parcel at 3.3 Hz. **I**. Spearman’s correlation across parcels between D_1_ receptor density and iPLV node centrality at 7.4 Hz. **J**. Spearman’s correlation across parcels between 5HT_1b_ receptor density and oCC node centrality at 3.3 Hz.

We next used a data-driven approach to identify clusters of frequencies that exhibited similar anatomical profiles of inter-areal connectivity patterns (Simola et al., 2022) in both PS and AC and then derived shared consensus frequency bands (Suppl. Fig. 2) for subsequent analyses: δ (1 – 3.5 Hz), low-θ (3.5 – 5.8 Hz), θ–α (5.8– 9.5 Hz), high-α (9.5 – 15 Hz), β (15 – 32 Hz), and γ (32 – 100 Hz). We estimated, as a measure of node centrality, in each frequency band, node strength per parcel as the mean iPLV or oCC, respectively, of this parcel with all other parcels (Suppl. Fig. 3A,B). Node centrality was positively correlated between iPLV and oCC at all frequencies (Fig. 2D), but more strongly in β and γ than in other frequency bands. To localize the differences, we obtained the z-score maps of node centrality and computed the difference PS - AC in each band (Suppl. Fig. 3C). The largest differences were observed around the occipital pole, which showed strong PS in all bands except β, but strong AC only in θ to α. Other anterior regions showed stronger AC than PS in most bands, while regions near the temporal pole and lateral sulcus showed higher PS in θ to α, but higher AC in δ and γ. These results corroborated the established notion that amplitude and phase coupling are overlapping, but distinct mechanisms in the human cortex.

### Frequency-band specific covariance of node centrality with receptor densities and principal components

We then set out to investigate the covariance of PS and AC node centrality with receptor and transporter densities and their principal components. We computed, for each single frequency (Suppl. Fig. 4) and each frequency band (Fig. 2 E-J), Spearman’s correlation coefficient between node strength and each receptor’s or transporter’s density across parcels. We first assessed the GABA_A_ and NMDA receptors, which through fast ionotropic transmission shape the generation of oscillations in local circuits of interacting glutamatergic excitatory pyramidal cells (PCs) and inhibitory GABAergic interneurons (INs) (Batista-Brito et al., 2018; Liley & Muthukumaraswamy, 2020; Lozano-Soldevilla et al., 2014; Lu et al., 2022; Middleton et al., 2008; Roopun, 2008). We found that both GABAa and NMDA receptor densities covaried positively with the PS node centrality in δ to α and γ bands, but with AC node centrality only in γ band (Fig. 3A,B, Suppl. Fig. 4). This result provides further evidence for the importance of these receptors for the generation of oscillations and also for the dissociation of PS and AC.

**Figure 3.**
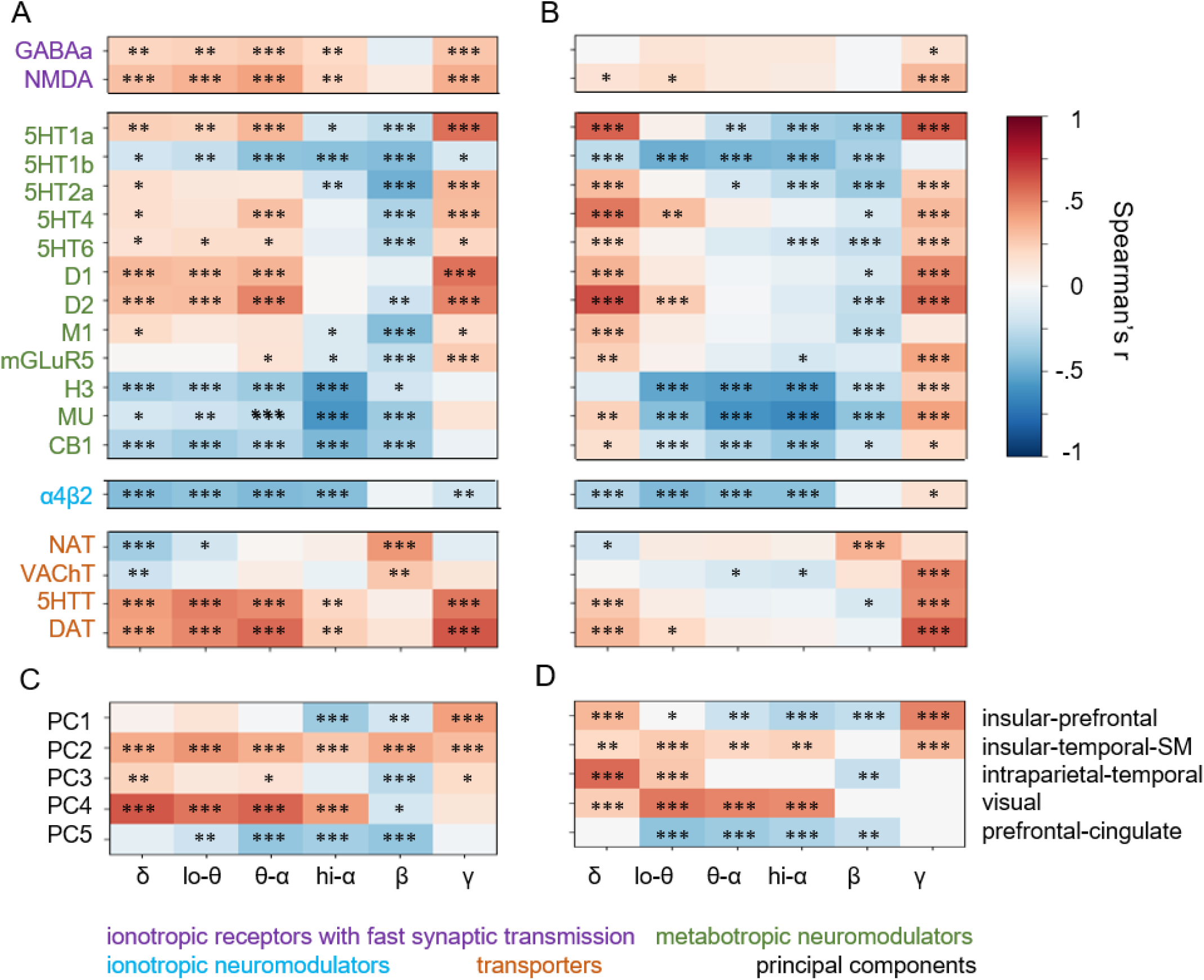
Covariances of density and node strength vary with frequency band and metric. **A**. Covariation of phase-coupling node centrality averaged within frequency bands with receptor and transporter density estimated across all cortical parcels with Spearman’s r (*: p < 0.05, **: p < 0.01, ***: p < 0.001). **B**. Same for amplitude-coupling node centrality. **C, D**. Same for covariation with principal components.**Abbreviations**: PC: principal component; 5HT: serotonin, 5HTT: serotonine transporter, D: dopamine, DAT: dopamine transporter, M1: muscarinic ACh receptor, mGLuR5: metabotropic (G-protein-coupled) glutamatergic receptor, NAT: noradrenaline transporter, VAChT: vesicular acetylcholine transporter, a4b2: α4ß2 nicotinic ACh receptor, H3: histamine receptor 3,, MU: mu-opioid receptor, CB1: cannabinoid receptor 1.

Among the receptors that act as neuromodulators on slower timescales and NT transporters, we observed similar patterns of covariance for all DA and 5HT receptors (except 5HT1b) and transporters as well as for M1 (muscarinic ACh) receptor and glutamatergic MGluR5. Similar to GABAa and NMDA, these generally covaried positively with node centrality of PS in lower bands and with that of both PS and AC in γ, but in addition also with AC node centrality in δ, and covaried negatively with both PS and AC node centrality in β band (meaning that peripheral rather than central nodes in these networks corresponded to high receptor density).

In contrast, 5HT1b, MU (mu-opioid), H_3_ (histaminergic), CB_1_(cannabinoid), and α_4_β_2_ (nicotinic ACh) receptors generally showed strong negative covariance with both PS and AC node centrality for δ to β bands, and positive covariance only with AC node centrality in γ band (and δ for MU and CB1). The noradrenaline and vesicular ACh transporters (NAT and AChT) showed negative covariance with node centrality in the lower frequencies, and some positive covariances in β and γ bands.

These results further underline the importance of neuromodulation by NTs such as 5HT, DA, and ACh via their effects on PC-IN circuitry (Batista-Brito et al., 2018; Börgers et al., 2008; Vinck et al., 2023). Notably, 5HT and DA tended to be highly expressed also in regions with strong resting-state γ node strength – which is in line with rodent studies showing modulatory effects of 5HT (Athilingam et al., 2017; Puig et al., 2010) and DA (L. Wang et al., 2020) on γ oscillations as well as computational modeling (D.-H. Wang & Wong-Lin, 2013). The lack of covariation between GABAa and NMDA densities and AC node centrality in the slower bands emphasizes that these two have different roles and operate on different timescales than neuromodulators such as DA and 5HT (Zhigalov et al., 2015).

These results of differences between covariance patterns of PS and AC with the receptor and transmitter densities support the view that PS and AC are related but dissociable, arising from partially distinct mechanisms, and support at least partially different functions (Engel et al., 2013; Siems & Siegel, 2020). In addition, also the the covariance of PS with receptor and transmitter densities d θ-α and high-α bands differed, which is in line with previous findings indicating that synchrony in low-α and high-α bands may serve different functions during task state (Lobier et al., 2018; Siebenhühner et al., 2016). The present findings suggest that differences between PS and AC as well as those between the frequency bands may arise in different cytoarchitectonic circuits.

DA, along with NA, has also been proposed to regulate long-range temporal correlations (LRTCs) that are the hallmark of brain critical dynamics, as well as the neuronal exhibition/inhibition (E/I) balance (Pfeffer, 2018). A recent study reported that the level of brain-wide inter-areal phase-connectivity and LRTCs are correlated and dependent on an individual’s E/I balance that acts as the main control parameter for critical dynamics (Fuscà et al., 2023). Therefore, our findings imply that NT receptor and transporter densities may influence synchronization and LRTCs within the framework of critical dynamics via the E/I balance.

Particularly noteworthy is that in the previous study of these receptor maps (Hansen et al., 2022) the α_4_β_2_, MU, CB_1_ and H_3_ receptors were found to possess the largest explanatory power with MEG signal power in δ, θ, α, and low-γ bands, while here, these were (along with 5HT1b) the receptors that showed the most negative covariation with node centrality. Notably, 5HT1b has been found to downregulate theta to beta oscillations in the frontal cortex of rodents (Kjaerby et al., 2016) and to inhibit other NTs such as DA, Glu, ACh, and GABA (Sari, 2004) and H3 has been implicated to affect cognitive function via regulation of NTs such as NA, DA, and 5HT (Esbenshade et al., 2008), while α4β2, MU, and CB1 are all known to be targets for drugs (nicotine, opioids, and cannabinoids) that strongly affect cognitive function.

In order to test for effects specific to biological sex, we repeated the covariance analysis separately for males (N=35) and females (N=32) and found that despite small variations, the larger covariance patterns were similar in both sexes (Suppl. Fig. 5).

### Principal components indicate neuroanatomical principles connecting receptor density to oscillatory connectivity

We then computed the covariance of the principal components with node centrality of PS and AC. We found that PC1 - which had positive loadings for all receptors - also showed both positive and negative covariation in a manner somewhat similar to that of the 5HT and DA receptors and transporters and especially that of mGluR5 which had the strongest loading. Covariations with both PS and AC NS was positive in γ bands and with PS also in θ-α band (although not in δ and low-θ), and negative especially with AC in the frequency bands in between (Fig. 3C-D). Somewhat similarly to PC1, PC3 covaried positively with PS in δ, θ-α and γ bands and with AC in δ and low-θ, and covaried negatively with both PS and AC in β. As PC1 notably represented the expression of receptors and transporters in frontal-temporal and medial regions, while PC3 reflected their expression (particularly 5HT1A, 5HT4, and D2) in temporal-occipital regions, these two may thus be seen as complementary kinds of neuromodulatory influence underlying the inter-areal connectivity in δ, θ, and γ bands.

In contrast to these two components, PC2 and PC4 showed positive covariance with NS across almost all frequency bands. PC2, which had highest loadings for DA and 5HT transporters and strongest expression in cingulate, temporal, and somatomotor regions, covaried positively with NS of both PS and AC in all frequency bands, in line with the cingulate cortex being a central hub in brain networks (Lavin et al., 2013; Leech et al., 2012) and DAT being important for working memory. PC4 was strongest in visual cortex and showed strong covariance with PS NS in all bands except β and with AC NS in θ and α bands, in line with well-known contributions of activity at NMDA and GABA receptors as well as lesser-known contributions of 5HT and DA receptors to the generation of low frequency oscillations in the visual cortex (Shaw et al., 2021).

Unlike other PCs, PC5 was negatively correlated with NS from low-θ to β bands. Interestingly, PC5 reflected densities of five receptors – 5HT1b, H3, α4β2, MU, and CB1 – showing mostly negative correlations with NS, especially in θ and α bands. Most of these had low expression in the lateral sulcus (except H3), but strong expression in the frontal cortex. PC5 thus may be seen as a common architecture for these receptors. These findings further support the notion that principal components capture underlying neuroanatomical principles and importantly indicate that these are related to inter-areal connectivity of neuronal oscillations.

### Limitations and outlook

It has been proposed that diversity of neuromodulators and their receptors in neuromodulatory ascending systems with high NT activity at specific receptors in particular regions enables circuit dynamics, supporting flexibility of behavior and cognition. In this study, we demonstrated the covariance of neurotransmitter receptor and transporter density with MEG-derived phase- and amplitude-based connectivity, i.e., the human dynome (Kopell et al., 2014). We specifically established how the heterogeneity in receptor and transmitter densities influences networks of phase synchrony and amplitude correlations. One limitation of the current study is the lack of usable density maps for some important receptors such as GABA_B_ that are known to be important for the generation of cortical oscillations in microcircuits (Liley & Muthukumaraswamy, 2020; Lozano-Soldevilla et al., 2014; Lu et al., 2022). Future studies will need to investigate how these microcircuits are influenced by the underlying neurotransmitter receptor and transmitter densities not only in resting state, but also during task (Boot et al., 2017; Cools, 2008; Puig & Gulledge, 2011).

The electromagnetic fields and currents underlying recorded MEG signals are thought to predominantly postsynaptic currents of colinearly oriented pyramidal neurons (Baillet, 2017; Hämäläinen et al., 1993), particularly those in infragranular layers (Siebenhühner et al., 2020), where α oscillations and synchrony dominate. As a general rule, receptor expression is largest in the upper layers (I-III), although with some exceptions for specific combination of region and receptor type (Palomero-Gallagher & Zilles, 2019). Future studies are needed to explore the layer-specific covariance between oscillatory connectivity and NT receptor and transporter maps.

Critically, understanding how NT receptor and transporter density distributions influence neuronal processing is essential for understanding the mechanisms of brain disorders. Abnormal oscillations and abnormal NT function characterize a wide range of brain disorders (Başar & Güntekin, 2008) including schizophrenia (Koch et al., 2016), depression (Celada et al., 2013; Fitzgerald & Watson, 2018; Nutt, 2008), ADHD (Banerjee & Nandagopal, 2015; Edden et al., 2012; Tellioglu & Robertson, 2001). Resolving the biological constraints on how oscillations are generated and modulated by neurotransmitters in a frequency- and region-specific manner (Batista-Brito et al., 2018; Roopun et al., 2010) may improve our understanding on the complex effects of pharmacological agents targeting neuromodulatory systems.

## Conclusion

The relationship between neurotransmitter activity on the one hand and oscillations and their connectivity on the other hand in the human brain is still far from understood. Here, we studied the covariance between NT receptor and transporter density distributions and node centrality in phase- and amplitude-based frequency-specific MEG-derived connectivity. Our findings demonstrate that networks of oscillatory connectivity are influenced by fundamental neuroarchitectonic principles underlying the distribution of NT receptors and transporters impacting connectivity in a region- and frequency-specific manner paralleling neuromodulatory pathways.

## Author contributions

F.S., S.P., and J.M.P conceptualized the study, F.S carried out data acquisition and all analyses. All authors wrote the manuscript.

## Acknowledgements

We thank Victoria Puig for helpful discussion and comments on an earlier version of this manusript.

## Methods

### MEG and MRI data acquisition

306-channel MEG (204 planar gradiometers and 102 magnetometers) was recorded from 67 healthy subjects (32 female, mean age 30.9 ± 8.3 years) with a Triux MEG (Elekta-Neuromag/MEGIN, Helsinki, Finland) at the BioMag Laboratory, HUS Medical Imaging Center. We recorded 10 minutes of eyes-open resting-state data from all participants. Bipolar horizontal and vertical EOG were recorded for the detection of ocular artifacts. MEG and EOG were recorded at a 1,000-Hz sampling rate.

T1-weighted anatomical MRI scans (MP-RAGE) were obtained for head models and cortical surface reconstruction at a resolution of 1 × 1 × 1 mm with a 1.5-Tesla MRI scanner (Siemens, Munich, Germany) at Helsinki University Central Hospital. Written informed consent was obtained from each subject prior to the experiment.

### Cortical parcellation and source modeling

Volumetric segmentation of MRI data, flattening, cortical parcellation, and neuroanatomical labeling were carried out using FreeSurfer software (http://surfer.nmr.mgh.harvard.edu). We used the original version of the Schaefer atlas with 200 parcels (Schaefer et al., 2018) based on the division of the brain into 17 functional subsystems (Thomas Yeo et al., 2011). MNE software (https://mne.tools/stable/index.html) (Gramfort et al., 2014) was used for the preparation of cortically constrained source models for MEG–MRI colocalization, forward and inverse operators. The source models had dipole orientations fixed to pial-surface normals and a 5-mm inter-dipole separation throughout the cortex, where hemispheres had between 5080–7645 active source vertices.

### MEG data preprocessing and filtering

Temporal signal space separation (tSSS) in the Maxfilter software (Elekta-Neuromag) was used to suppress extracranial noise from MEG sensors and to interpolate bad channels. We used independent components analysis (ICA) adapted from the MATLAB toolbox Fieldtrip (http://www.fieldtriptoolbox.org) to extract and identify components that were correlated with ocular artifacts (identified using the EOG signal), heartbeat artifacts (identified using the magnetometer signal as a reference), or muscle artifacts. After artifact exclusion, the time-series data were filtered into NB time series using a bank of 53 Morlet filters with wavelet width parameter m = 5 and approximately log-linear spacing of center frequencies ranging from 1.1 to 315 Hz.

We estimated vertex fidelity to obtain fidelity-weighted inverse operators that improve reconstruction accuracy, as described in (Siebenhühner et al., 2020). For connectivity analysis, we further estimated parcel fidelity and cross-parcel mixing and removed parcels with fidelity < 0.1 and parcel pairs with cross-parcel mixing PLV > 0.4, for which the likelihood of spurious connectivity was notably increased. In total, 12.7% of edges were removed.

### Analysis of interareal phase synchronization and cross-frequency connectivity

To estimate pairwise synchronization between parcels, we estimated the complex phase-locking value (cPLV):

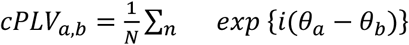

where *θ*_*a*_ and *θ*_*b*_ are the instantaneous phase time series of the complex analytical narrowband time series *X*_*a*_ and *X*_*b*_, and *N* is the number of samples *n*.

From the cPLV, the “classical” PLV can be derived as *PLV* = |*cPLV*| and the imaginary *PLV* as *iPLV* = |*im*(*cPLV*)|. The imaginary PLV is insensitive to zero-lag false-positive interactions which are often spurious due to residual linear mixing after inverse modeling (J. M. Palva et al., 2018).

Amplitude correlations were computed between the amplitude envelopes of the narrowband time series *X_a_* and *X_b_* with the correlation coefficient (CC) and the orthogonalized correlation coefficient (oCC). In the latter, the time series X_b_ is orthogonalized with respect to *X_a_*, which also has the effect of eliminating spurious connections (Brookes et al., 2011; Hipp et al., 2012).

We then computed the spatial similarity of node strength matrices between frequencies and then used the Louvain algorithm (Blondel et al., 2008) to identify frequency clusters, for both iPLV and oCC separately and then derived from these consensus ranges.

### Neurotransmitter receptor and transporter maps and correlation with node strength

We downloaded the code and dataset from https://github.com/netneurolab/hansen_receptors and computed the receptor density in each of the 200 parcels for 35 different receptor and transporter maps. This dataset has been recently described in detail in (Hansen et al., 2022). Where several maps were available for the same receptor or transporter, we computed the mean value, with each map weighted by the number of subjects, arriving at 19 density maps. Correlation with mean node strength per parcel was computed for each map in each individual frequency using Spearman’s correlation coefficient, as well as in frequency bands (with node strength having been averaged over individual frequencies) across all 200 parcels.

### Principal component analysis

We used the PCA algorithm from the python toolkit scikit-learn to identify the first 4 principal components (PCs) underlying the receptor and transporter maps. We then computed the strength of each PC in functional Yeo subsystems by summing PC strength over all parcels belonging to a subsystem. We further estimated the correlations of each component with individual receptors as well as with PS and AC in the six frequency bands using Spearman’s correlation.

## Supplementary materials

**supp. Figure 1.**
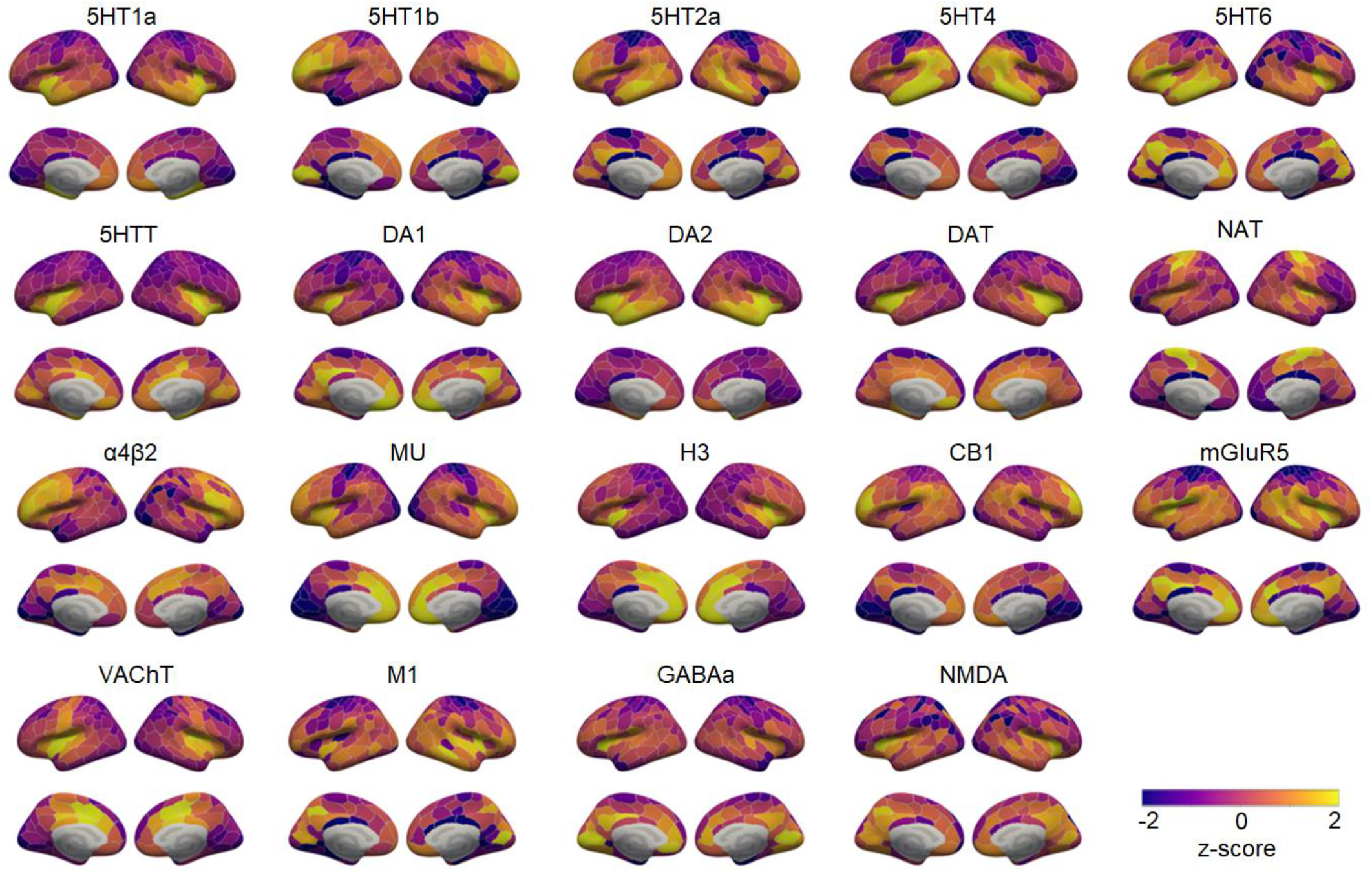
Receptor density maps, z-scored.

**supp. Figure 2.**
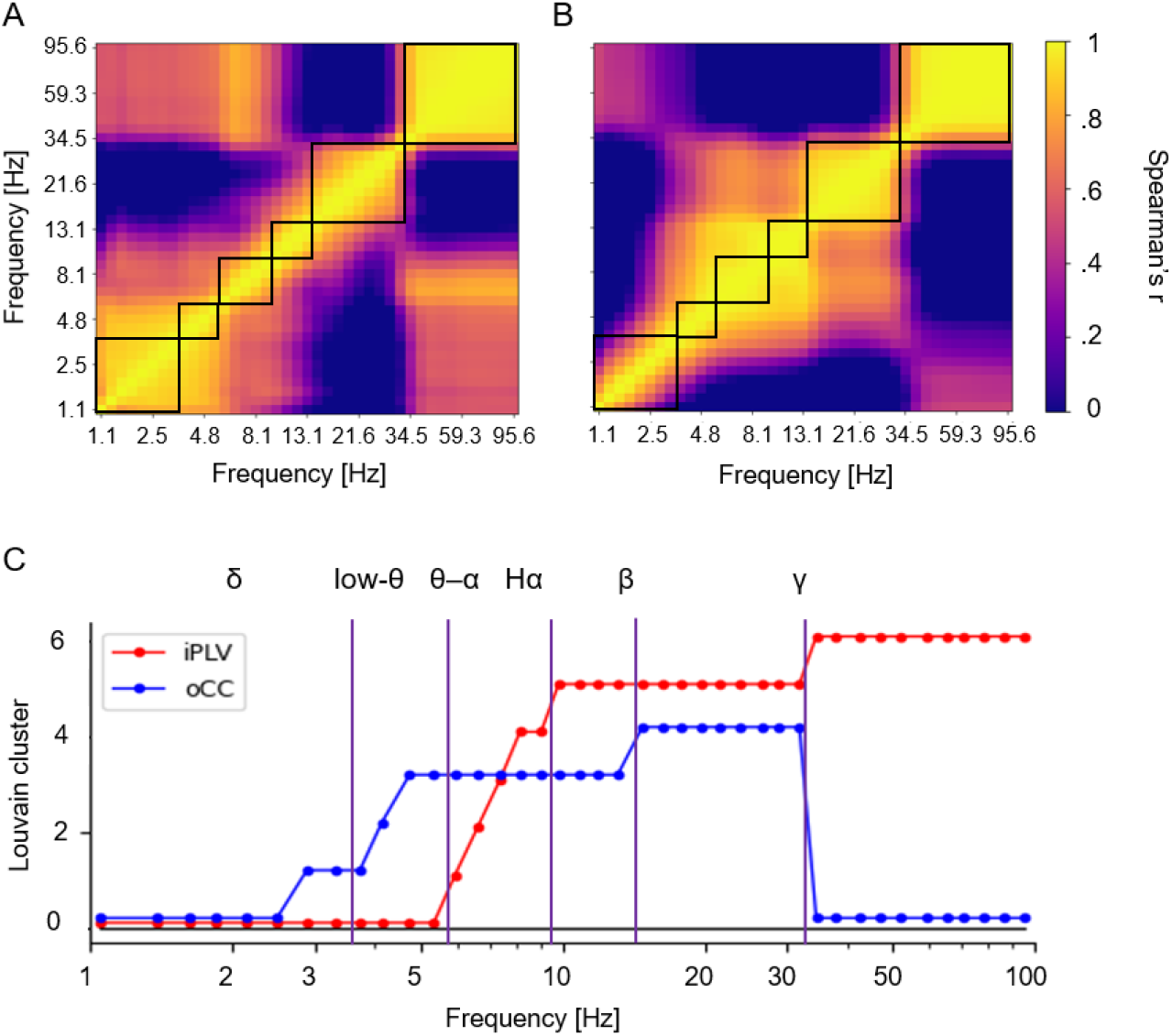
**A,B**. Pairwise similarity of node strength between frequencies for iPLV and oCC. **C**. Frequency clusters obtained with the Louvain algorithm and consensus frequency bands.

**supp. Figure 3.**
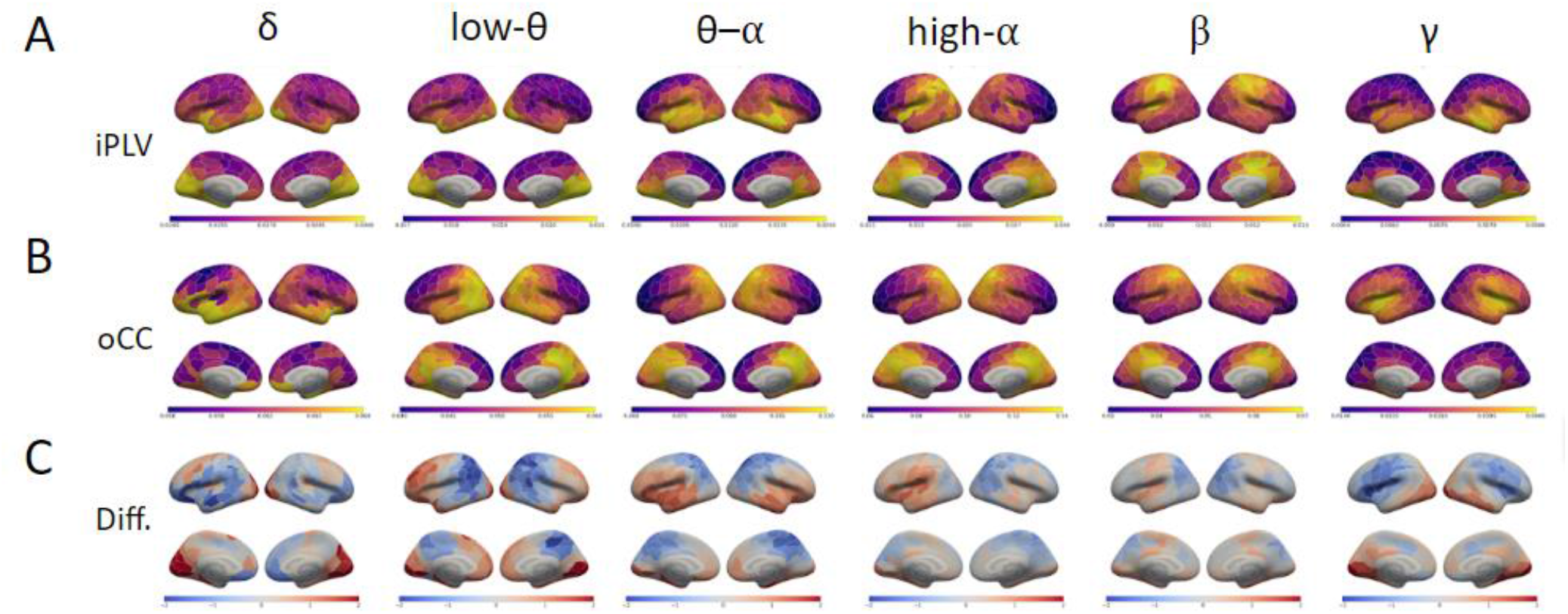
**A**. Mean iPLV node strength per cortical parcel and frequency band. **B**. Same for oCC. **C**. Difference of z-scored PS node strength minus z-scored AC node strength.

**supp. Figure 4.**
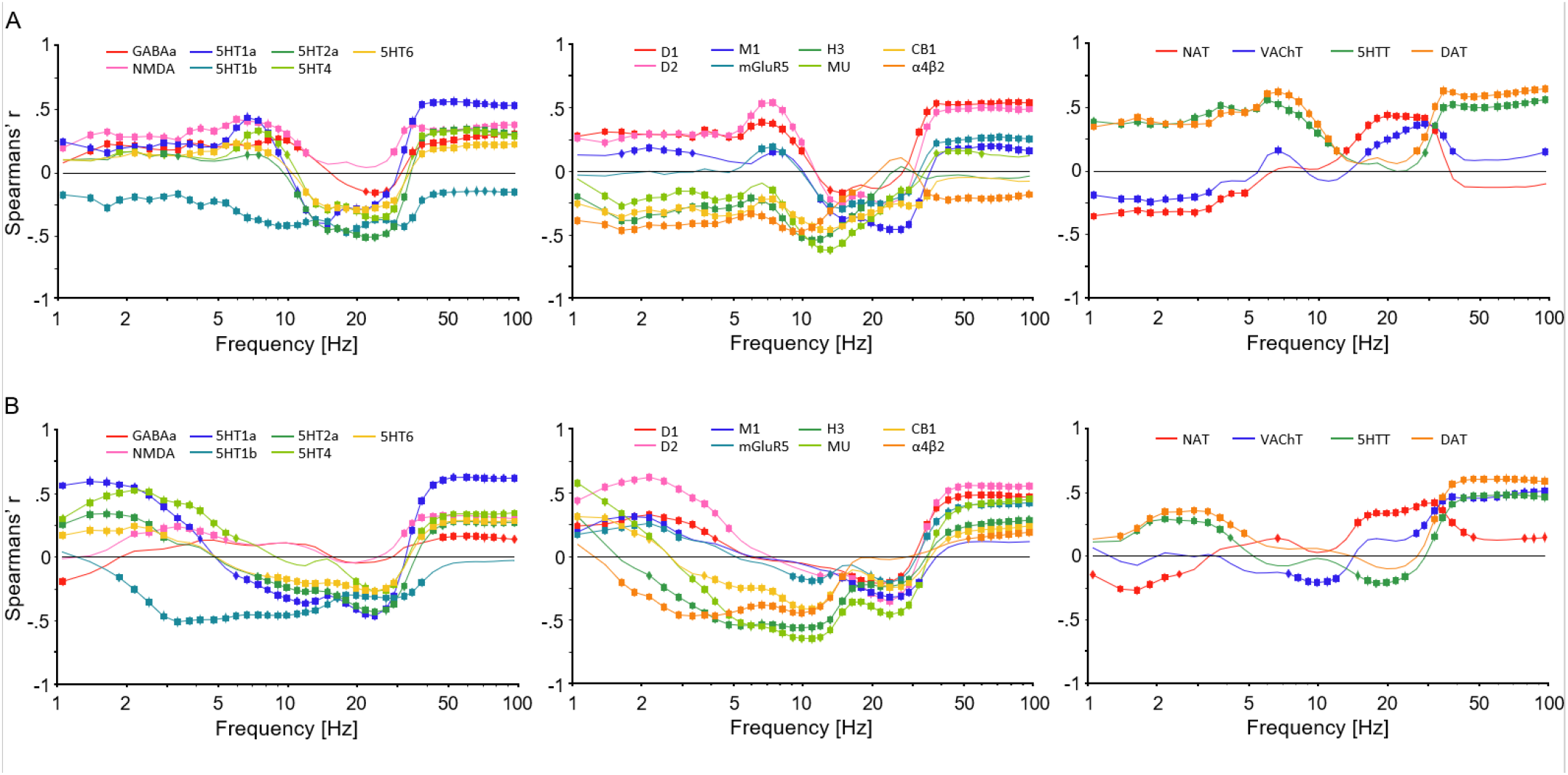
Covariance between node strength and receptor density estimated across cortical parcels, as in Figure 3, but for individual narrow frequencies. Significant correlations are marked with diamonds (Spearman’s r, p < 0.05) and those that remained significant after correction for multiple comparisons with Benjamini-Hochberg method with squares.

**supp. Figure 5.**
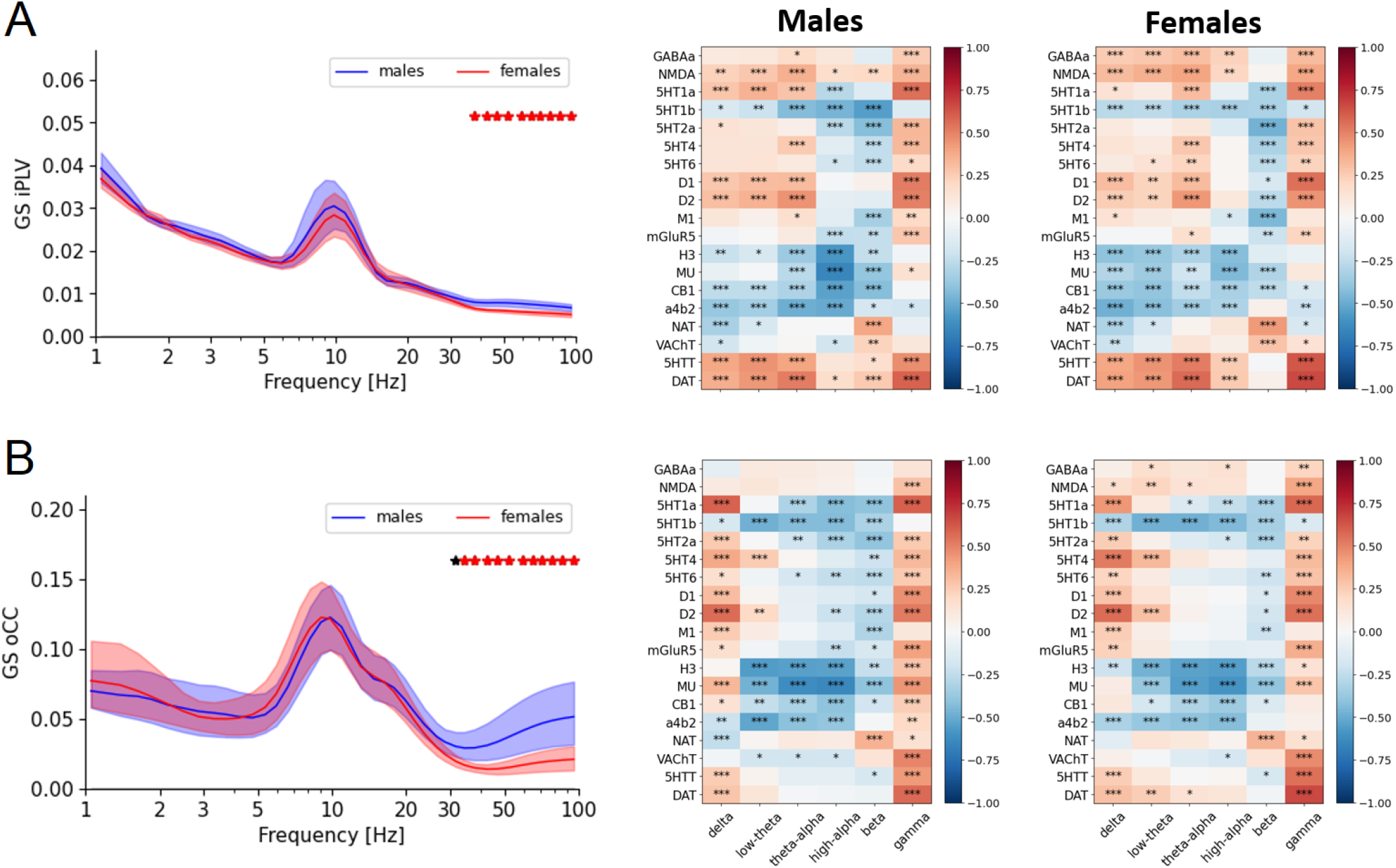
**A**. Left: Mean iPLV graph strength for male and female subjects. Asterisks indicate significant group differences (Mann-Whitney-U test, p < 0.05, red if significant after correction with Benjamini-Hochberg). Right: Covariance of node centrality with neurotransmitter receptor and transporter density maps (right), as in Figure 3. **B**. Same for oCC.

